# Loss of adipocyte identity through synergistic repression of PPARγ by TGF-β and mechanical stress

**DOI:** 10.1101/604231

**Authors:** Ewa Bielczyk-Maczyńska, Brooks Taylor, Cayla M Miller, Michael L Zhao, Arnav Shah, Zahra Bahrami-Nejad, Alexander R Dunn, Mary N Teruel

**Affiliations:** Department of Chemical & Systems Biology, Stanford University, Stanford, CA 94305, USA; Department of Chemical Engineering, Stanford University, Stanford, CA 94305, USA

## Abstract

Adipocytes convert into myofibroblasts in a TGF-β-dependent mouse model of fibrosis. The molecular steps and timing underlying this conversion are poorly understood, hindering development of antifibrotic therapies. Here we used two single-cell approaches, lineage tracing and live-cell imaging of an adipocyte marker PPARγ, to track the fate of adipocytes induced to convert by TGF-β. We found that TGF-β alone was not sufficient to activate the TGF-β pathway and to induce myofibroblast conversion in cells with high PPARγ expression. However, robust conversion was observed when an additional PPARγ-inhibiting stimulus, mechanical stress applied by increasing adhesion area on a stiff matrix, was applied simultaneously with TGF-β. We show that the PPARγ downregulation in response to increased adhesion area required both fibronectin and a sufficiently stiff extracellular matrix (ECM) and was partially mediated by Rho. Our results show for the first time the order of the molecular processes driving fat tissue fibrosis and the requirement for signal convergence for the loss of adipocyte identity.

## Introduction

Cell plasticity, the ability of differentiated cells to convert into other cell types, underlies pathogenesis of many diseases including diabetes^1^ and cancer^2^. Plasticity of adipocytes (fat cells) includes a reversible dedifferentiation into proliferative adipocyte progenitors in the mammary gland during lactation^3^ and a pathogenic conversion into myofibroblasts in dermal fibrosis^4^. Fibrosis is driven by the accumulation of myofibroblasts, which can be derived from a variety of cellular sources, including tissue-resident fibroblasts, adipocyte progenitors^5,6^ and adipocytes^4^. Normally myofibroblasts appear after tissue injury in response to local profibrotic signals such as the TGF-β cytokine, facilitate the wound healing process, and subsequently undergo apoptosis. However, in fibrosis the presence of myofibroblasts becomes permanent, and they produce excess extracellular matrix (ECM) leading to increased tissue stiffness, which in itself may be one of the drivers of the disease progression^7^. The current lack of antifibrotic therapies underscores the necessity to understand the mechanisms of fibrosis development and progression.

While the molecular networks regulating adipocyte differentiation have been well studied^8^, the mechanisms of adipocyte identity loss remain incompletely understood. Here we present two complementary methods that allow for quantitative tracking of adipocyte identity loss at the single-cell level. PPARγ, a transcription factor which has been shown to be sufficient and required to drive adipocyte differentiation^9, 10^, is critical for maintaining adipocyte function^11,12^. We show that cells with high expression of PPARγ inhibit signaling by the profibrotic cytokine TGF-β. However, TGF-β stimuli lead to PPARγ downregulation when they are applied to cells simultaneously with adhesion-induced mechanical stress on a stiff fibronectin-containing ECM. Together, these findings establish that an integration of chemical and mechanical stimuli occurring in a specific order drives fibrosis progression in fat tissue. These results also suggest that mechanotransduction pathways, and in particular Rho kinase-dependent signaling, could become novel molecular targets for scleroderma therapies.

## Results

### Adipocyte-myofibroblast conversion is detected at single-cell level by lineage tracing

To detect the adipocyte-myofibroblast switch *in vitro*, we used cells derived from transgenic Adipoq:Cre mT/mG mice in which the adipocyte-specific adiponectin promoter drives the expression of Cre recombinase, causing an irreversible switch from membrane red (Tomato) to membrane green (GFP) fluorescence in adipocytes (Fig. 1a). Adiponectin has been shown as the most reliable marker for lineage tracing of mature adipocytes because its expression does not label fat progenitors or other non-adipogenic cell populations present in the fat pad^13^. To test if the adipocyte-myofibroblast conversion occurs in Adipoq:Cre mT/mG primary adipocytes, we first isolated the preadipocyte-containing stromal vascular fraction (SVF) from subcutaneous fat pads and subjected these cells to an adipogenic differentiation protocol *in vitro*. Given that mechanical cues are known to drive myofibroblast differentiation from fibroblasts^14^, we decided to use mechanical cues to try to induce myofibroblast differentiation of the primary SVF-derived adipocytes. We thus passaged the SVF-derived adipocytes at subconfluence which increases the area of cell interaction with the stiff culture plate^15^. Simultaneously, cells were stimulated with the profibrotic cytokine TGF-β or not. Marker gene expression was analyzed using immunofluorescence (IF) staining at several timepoints (Fig. 1b). Based on the irreversible GFP expression in the SVF-derived adipocytes, we could assess different cell states assumed by these cells at later time points, using co-staining of myofibroblast (α-SMA) and adipocyte (PPARγ, C\EBPα) markers (Fig. 1c). We observed that while GFP-positive cells often had residual Tomato fluorescence (Fig. 1d-e), high GFP expression could be reliably used to detect the adipocyte-derived cell population (Fig. 1f). To detect myofibroblasts within this GFP-positive population, we used the presence of α-SMA-containing stress fibers as the most robust marker of myofibroblasts^14^(Fig. 1d, Supplementary Fig. S1). Conversely, the expression of PPARγ (Fig. 1e) and C\EBPα in GFP-positive cells was used to detect cells which remained in the adipocyte state. Under TGF-β treatment the number of GFP-positive myofibroblasts significantly increased over six days (Fig. 1g). This effect was most likely attributable to the increased number of GFP-positive cells surviving under TGF-β compared to the control condition (Supplementary Fig. S2). Adipocyte marker proteins PPARγ and C\EBPα were downregulated in the GFP-positive population after replating, and treatment with TGF-β exacerbated this adipocyte marker loss, leading to the absence of PPARγ- and C\EBPα-positive cells in the GFP-positive population after six days (Fig. 1h).

**Figure 1.**
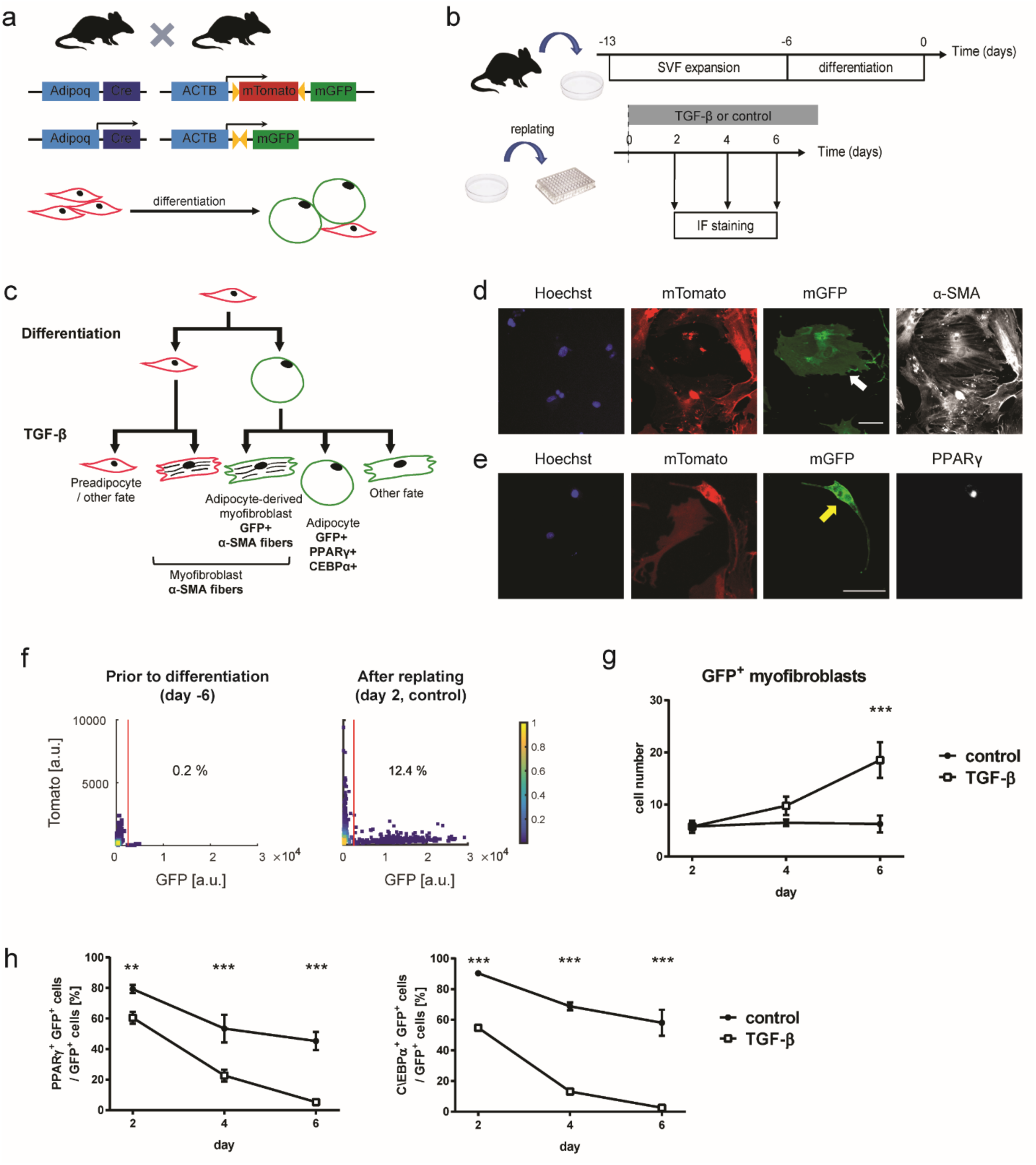
Lineage tracing allows for measurement of the dynamics of the adipocyte-myofibroblast transition and shows that TGF-β induces the conversion of primary adipocytes into myofibroblasts. **a** In the transgenic mouse system Adipoq:Cre mT/mG, preadipocytes express fluorescent protein tdTomato at the plasma membrane. The mTomato (RFP) irreversibly switches to mGFP under the control of an adipocyte-specific adiponectin (Adipoq) promoter. **b** Primary SVF cells were expanded and differentiated into adipocytes *in vitro*. The differentiated cells were then replated at the start of the experiment in order to introduce a mechanical stimulus resulting from the process of adhesion and spreading on a stiff substrate. TGF-β was added to the media immediately after replating and cells were analyzed at day 2, 4 or 6 using IF staining. **c** Lineage tree showing possible cell states after TGF-β treatment. Lineage markers used to quantify number of cells in each category shown in bold. **d-h** Primary SVF was expanded and differentiatied *in vitro*, replated, treated with **d** TGF-β or **e** control media for six days, and subjected to IF staining against **d** α-SMA or **e** PPARG. **d** An adipocyte that converted to the myofibroblast phenotype is denoted by a white arrow. **e** An adipocyte which maintained PPARγ expression is denoted by a yellow arrow. Scale bar: 100 µm. **f** Single-cell distribution of Tomato and GFP expression in primary SVF prior to differentiation (day -6) and 2 days after replating in control media. Cut-off used to filter GFP-positive cells at day two and percentage of GFP-positive cells are indicated. **g** Number of GFP-positive myofibroblasts per well over time. **h** Percentage of GFP-positive cells which express adipocyte markers PPARγ and C/EBPα. **g-h** Results of one experiment representative for two independent experiments. Two-tailed Student *t* tests with Benjamini-Hochberg correction; FDR=0.01; n=4 technical replicates, **, p<0.01; ***, p<0.001. GFP-positive cells/replicate/time point > 32.

### Passage on a stiff substrate is required for the TGF-β-induced downregulation of adipocyte marker expression

Because of the observed downregulation of adipocyte marker expression after cell replating, we decided to test whether the mechanical stimulus introduced by replating was required for the TGF-β-induced downregulation of PPARγ and C\EBPα. To this end, we investigated whether adipocyte markers are downregulated by TGF-β in primary adipocytes which were not replated at subconfluence. In the Adipoq:Cre mT/mG model the expression of adipocyte lineage marker GFP in the cell membrane led to a high incidence of mis-assignment of GFP-negative cells as GFP-positive in a confluent population of cells. In order to faithfully assign confluent cells to adipocyte-(GFP-positive) and non-adipocyte-derived (GFP-negative) populations, we turned to a lineage tracing mouse model with nuclear localization of GFP/Tomato (Adipoq:Cre nT/nG mice, Fig. 2a). Analogously to the approach presented in Figure 1, we differentiated the SVF cells *in vitro*. However, cells were subjected to TGF-β or control treatment without passaging (Fig. 2b). Immunofluorescence staining revealed that virtually all GFP-positive cells maintained their high PPARγ and C\EBPα expression through six days of analysis, irrespective of TGF-β presence (Fig. 2c-d). Altogether, this finding suggests an interplay between profibrotic cytokine TGF-β and mechanical stimuli in driving a cell identity change in adipocytes.

**Figure 2.**
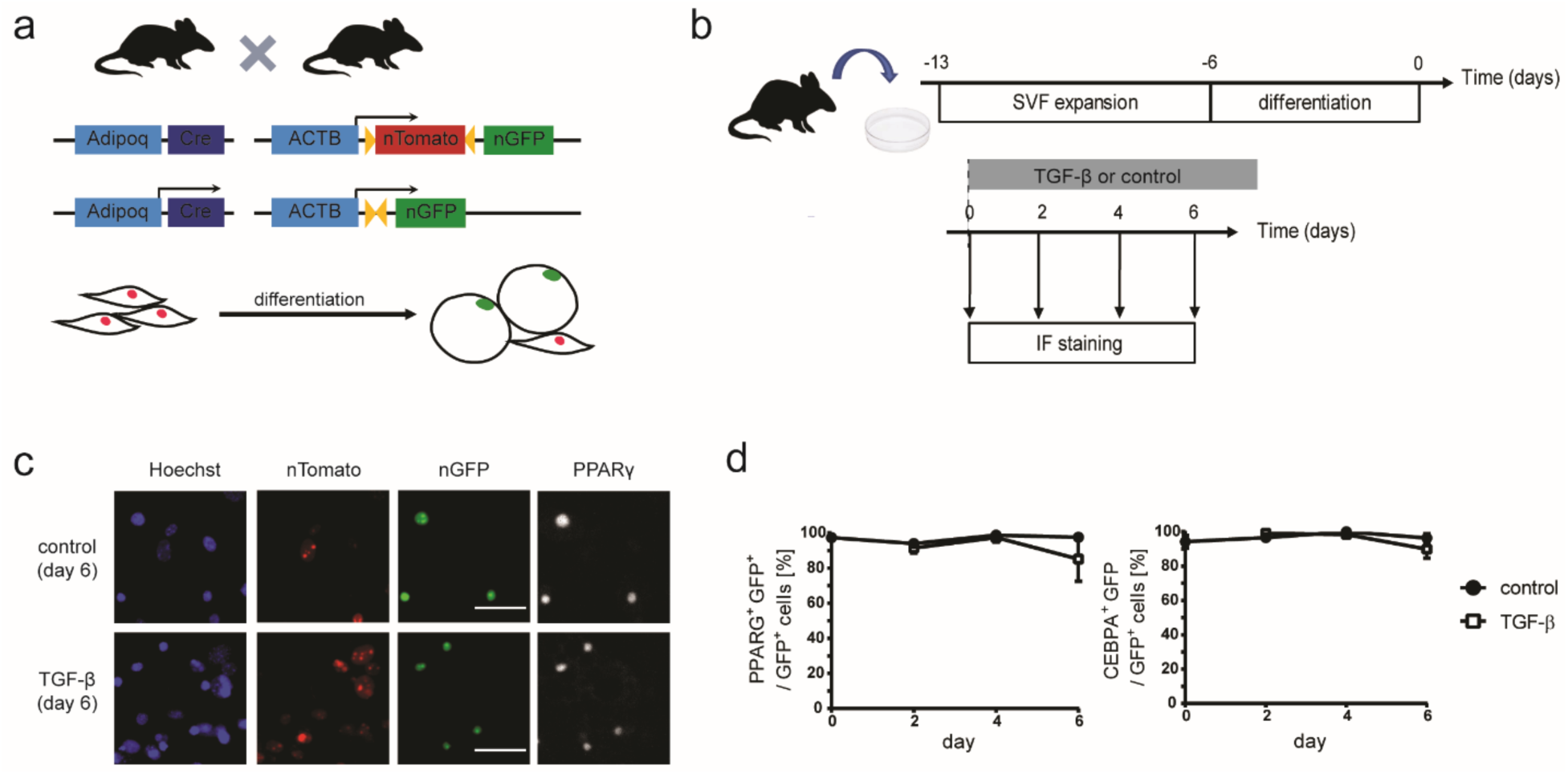
Lack of a mechanical stimulus associated with cell passage onto a stiff substrate prevents TGF-β-induced downregulation of adipocyte marker proteins. **a** In the transgenic mouse system used, preadipocytes express the fluorescent protein tdTomato in cell nucleus. The nuclear tdTomato (nTomato) irreversibly switches to nuclear GFP (nGFP) under the control of the adipocyte-specific adiponectin (Adipoq) promoter. **b** Primary SVF cells were expanded and differentiated into adipocytes *in vitro*. TGF-β was added to the media at the end of differentiation (day 0) and cells were analyzed at days 0, 2, 4 or 6 using IF staining. **c** Representative fluorescent images of IF staining against PPARγ at day 6 after adding stimulus. Scale bar: 50 µm. **d** Percentage of GFP-positive cells which expressed adipocyte markers PPARγ and C/EBPα. Two-tailed Student *t* tests with Benjamini-Hochberg correction; FDR=0.01; n=3-8 technical replicates, all time points p>0.05.

### PPARγ downregulation by TGF-β is dependent on ECM stiffness and composition

Given the observed interplay between mechanical stimuli and TGF-β, we decided to investigate whether TGF-β affects the mechanical interactions between adipocytes and their microenvironment. To this end, we measured cell area as a proxy for interaction with the microenvironment in differentiated Adipoq:Cre mT/mG SVF cells. After replating and at 48 hours of treatment with TGF-β or control media, we used immunofluorescence staining of GFP and PPARγ in the same cells to quantify the single-cell spreading area (Fig. 3a). TGF-β treatment caused a strong increase in the percentage of large-area cells within the GFP-positive population (Fig. 3b). Additionally, both in the control sample and for cells treated with TGF-β, large cell area was associated with low PPARγ expression (Fig. 3c). Based on these findings, we hypothesized that TGF-β leads to an increase in the mechanical interaction of adipocytes with their microenvironment.

**Figure 3.**
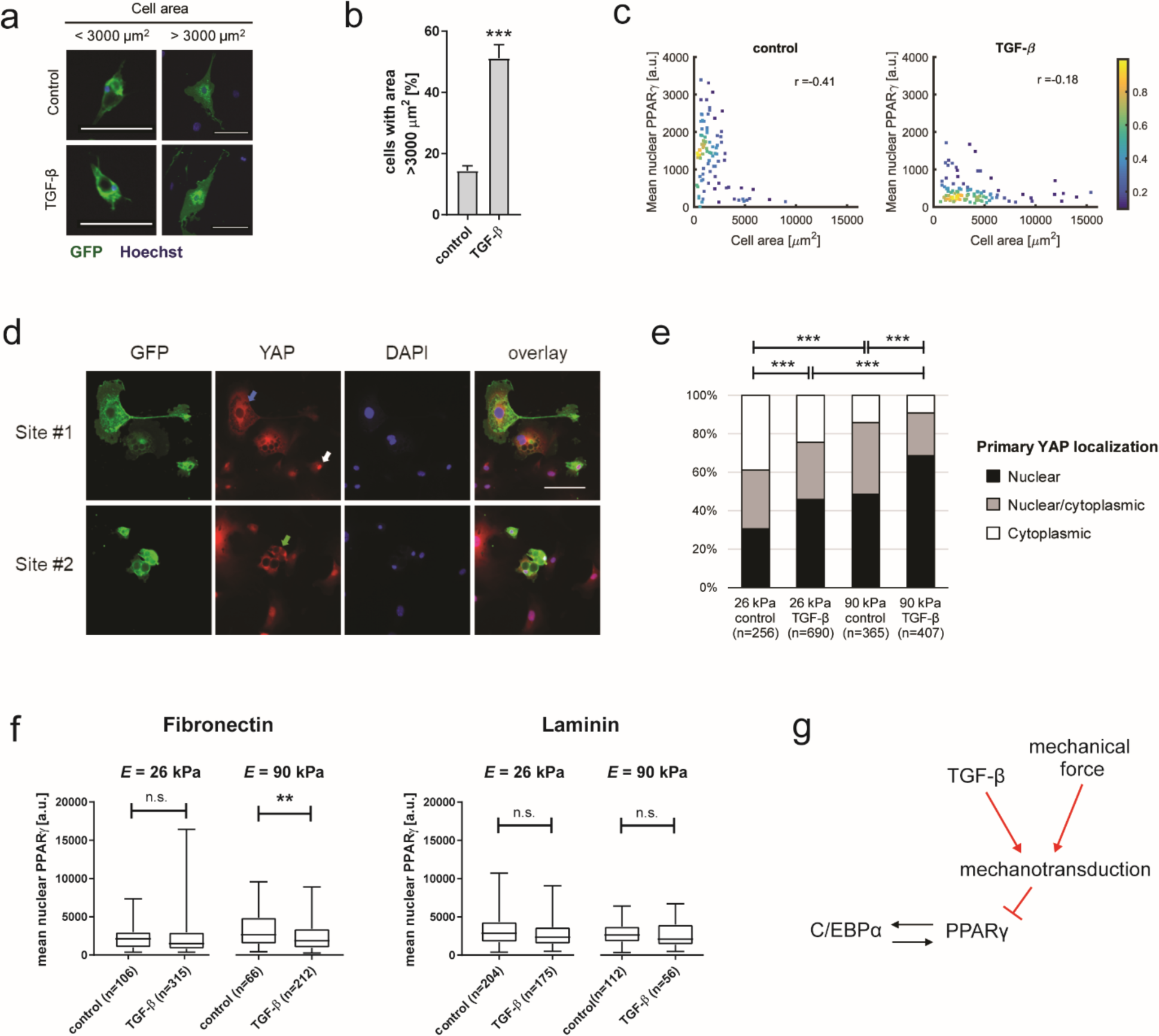
PPARγ downregulation by TGF-β in SVF-derived adipocytes is dependent on ECM stiffness and protein composition. **a** Representative images of GFP-positive (green) SVF-derived adipocytes replated on tissue culture plastic and fixed after 48h. Nuclei were counterstained with Hoechst (blue). Scale bar: 100 µm. **b** Quantification of GFP-positive cells with large (> 3,000 µm^2^) cell area in control and TGF-β-treated conditions. One-tailed Student t-test; ***, p<0.001; n=4 technical replicates. **c** GFP-positive SVF-derived adipocytes were subjected to IF staining against PPARγ and GFP at 48h after plating, and mean nuclear PPARγ expression and cell area were quantified. r, Pearson correlation coefficient. Color indicates local density of data points. **a-c** Results of one experiment representative for two independent experiments. **d** Images showing YAP localization in SVF-derived adipocytes plated on a polyacrylamide gel coated with fibronectin (Young’s Modulus, *E*=90 kPa). Representative cells with predominantly nuclear (white arrow), cytoplasmic (blue arrow) and no predominant localization (green arrow) of YAP are indicated. Scale bar: 100 µm. **e** Quantification of cellular localization of YAP in SVF-derived GFP-positive cells plated on fibronectin-coated polyacrylamide gels (*E*=26 kPa or 90 kPa) in the presence of TGF-β and in control conditions. Chi-squared test with Bonferroni correction; ***, p<0.001. Results of one experiment representative for two independent experiments. **f** Quantification of PPARγ intensity in SVF-derived GFP-positive adipocytes plated on polyacrylamide gels coated with fibronectin or laminin, of Young’s modulus 90kPa or 26 kPa. Cells were treated with either TGF-β or control media and analyzed after 24h. Number of analyzed GFP-positive cells per group are shown. One-way ANOVA with Sidak’s multiple comparisons test; **, p<0.01. **g** Schematic summarizingthe data in Figure 3.

### Nuclear localization of YAP is increased both by high ECM stiffness and by TGF-β

To further explore the possible activation of mechanotransduction pathways in adipocytes, we measured the nuclear translocation of Yes-associated protein (YAP), a transcriptional coactivator whose localization to the nucleus often correlates with ECM rigidity^16^. We plated differentiated SVF cells from Adipoq:Cre mT/mG mice on polyacrylamide gels of different stiffnesses, treated them with TGF-β or control media, and assessed the cellular localization of YAP in GFP-positive cells after 24 hours using IF staining (Fig. 3d). The occurrence of nuclear YAP localization in GFP-positive cells was increased both by higher substrate stiffness and by TGF-β stimulation, and these effects appeared additive (Fig. 3e), suggesting that TGF-β increased localization of YAP to the nucleus independently of ECM rigidity. Finally, to test whether the effect of TGF-β on PPARγ downregulation was modulated by ECM rigidity, we seeded differentiated primary SVF from Adipoq:Cre mT/mG mice on polyacrylamide gels of different stiffnesses, treated them with TGF-β or control media, and assessed the level of PPARγ in GFP-positive cells 24 hours later using IF staining. TGF-β caused a significant decrease in PPARγ expression level on the stiffer gel (Young’s modulus *E*=90 kPa vs. 26 kPa), but only when it was coated with fibronectin and not with laminin (Fig. 3f). We concluded that TGF-β can cause PPARγ downregulation in adipocytes when the ECM stiffness and composition are permissive (Fig. 3g).

### TGF-β stimulation and a mechanical stimulus introduced by passaging onto a stiff substrate act simultaneously to irreversibly downregulate PPARγ

To obtain deeper understanding of the dynamic interplay between TGF-β and mechanical stimuli in the loss of adipocyte state, we took advantage of another cell model, the adipogenic mouse cell line OP9^17^. This cell line has been previously used by us to obtain insight into the dynamics of adipocyte differentiation^18, 19^. First, to validate that TGF-β exerts profibrotic changes in this system, we subjected differentiated OP9 cells to TGF-β treatment for 96h and analyzed bulk gene expression changes at the mRNA level. As expected, TGF-β caused upregulation of myofibroblast markers *Acta 2* and *Col1a1*, as well as downregulation of adipocyte markers *Pparg, Cebpa* and *Fabp4*. In addition, we did not observe upregulation of preadipocyte markers *Dlk1* and *Pdgfra*, suggesting that under TGF-β treatment only the myofibroblast population expanded over time (Fig. 4a). Interestingly, in contrast to primary differentiated SVF cells, in OP9 cells TGF-β treatment led to the formation of cell clumps, which could be a source of a mechanical stimulus in this system (Supplementary Fig. S3).

**Figure 4.**
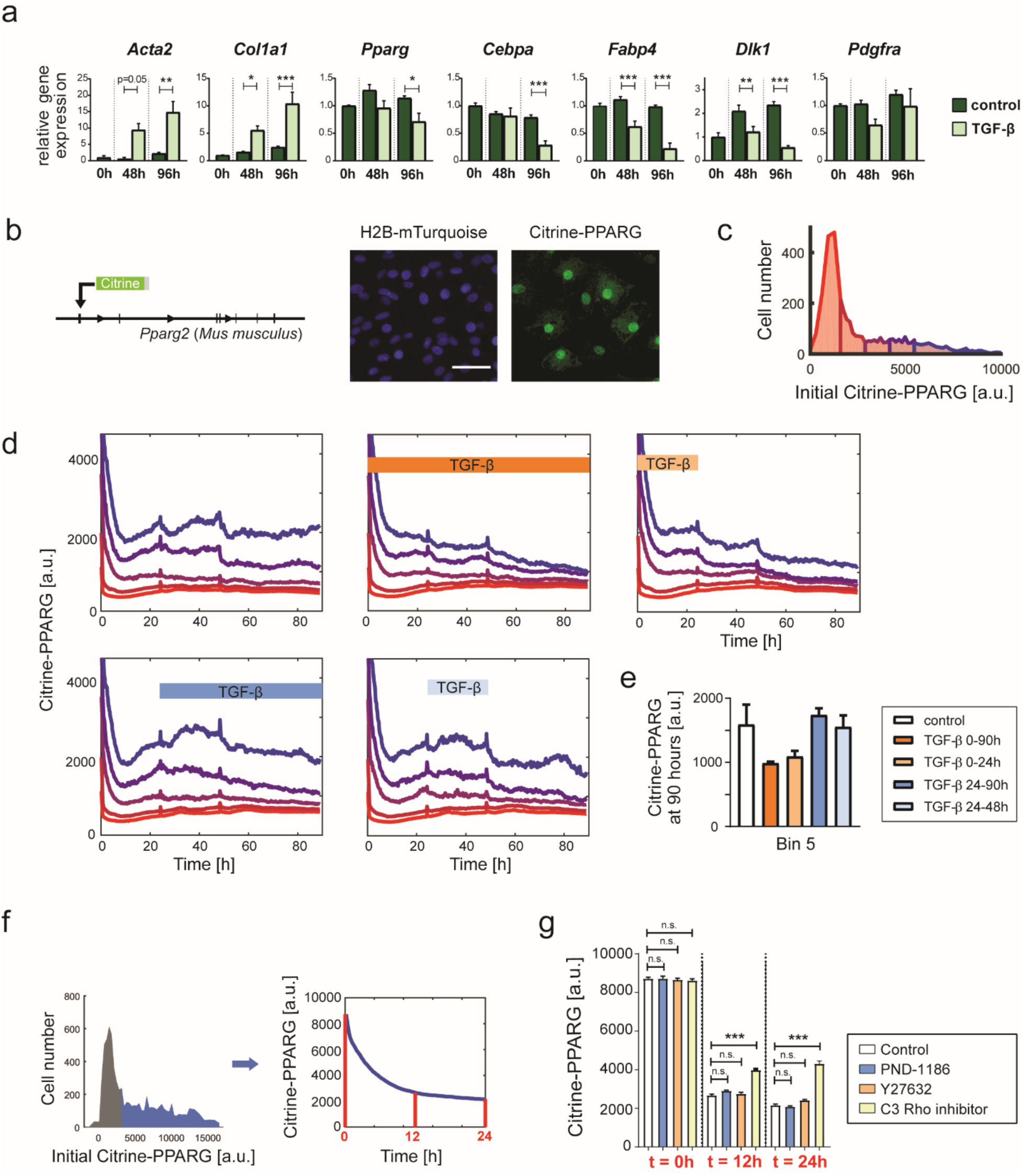
PPARγ downregulation in adipocytes requires synergistic timing of a mechanical stimulus introduced by passaging onto a stiff substrate and TGF-β treatment. **a** TGF-β induces upregulation of myofibroblast markers with concomitant downregulation of adipocyte and preadipocyte markers. TGF-β or control media added to OP9 cells at the end of differentiation (0h). RT-qPCR, n=4 technical replicates. One-way ANOVA with Sidak’s test. *, p<0.05; **, p<0.01; ***, p<0.001. Results of one experiment representative for two independent experiments. **b** Endogenous *Pparg2* locus in mouse OP9 cells was tagged with mCitrine (YFP) using CRISPR-mediated genome editing^19^. Fluorescent nuclear marker H2B-mTurquoise was added. Schematic of CRISPR-based tagging and representative images of differentiated cells shown. Scale bar, 100 um. **c** Distribution of single-cell mCitrine-PPARG expression at the end of differentiation protocol. **c-e** Cells were binned and the bins were color-coded based on initial mCitrine-PPARG expression. **d** mCitrine-PPARG downregulation occurs if TGF-β stimulation happens during the first 24 hours when cells are adhering after replating. Windows of treatment with 2 ng/ml TGF-β are shown. Plots show median traces for each bin. Culture media was replaced at 24h and 48h for all conditions. **e** Quantification of median endpoint (90h) mCitrine-PPARG expression in Bin 5 (the bin with the highest initial expression). n=4 technical replicates, all p > 0.05. **f** Workflow used to identify pathways involved in PPARγ downregulation in adipocytes following replating. Initial mCitrine-PPARG expression was used to identify differentiated (mCitrine-PPARG-high, marked in blue) cells. The mean mCitrine-PPARG expression in these cells was compared at 0, 12 and 24h after replating the cells with immediate stimulus addition. **g** Replated, differentiated OP9 cells were treated with chemical inhibitors of FAK (PND-1186), ROCK (Y27632), or Rho (C3 Rho inhibitor I) to test for rescuing effect on mCitrine-PPARG downregulation. The inhibitors were added at replating. n=4 technical replicates, >346 cells per replicate. One-way ANOVA with Sidak correction; **, p<0.01; ***, p<0.001; n.s. – not significant. Results of one experiment representative for 2-3 independent experiments per inhibitor.

Next, we took advantage of an OP9 cell line in which the key adipocyte marker PPARγ2 has been endogenously tagged with a fluorescent protein mCitrine (mCitrine-PPARG line)^19^. Additionally, this line contains a fluorescent nuclear marker H2B-mTurquoise (Fig. 4b). This model allows for simultaneous tracking of endogenous PPARγ levels in thousands of differentiated cells. At the end of a standard four-day differentiation protocol, cells show a range of individual mCitrine-PPARG expression levels (Fig. 4c), with high mCitrine-PPARG present in differentiated adipocytes^19^. We used live tracking of differentiated mCitrine-PPARG cells, grouping the cells by their mCitrine-PPARG expression level at the beginning of live imaging (Fig. 4c). To address whether a specific order of a mechanical stimulus and TGF-β stimulation was required for adipocyte identity loss, differentiated mCitrine-PPARG OP9 cells were passaged at subconfluence and tracked for 90 hours using live-cell fluorescent imaging. Cells were additionally stimulated with 2 ng/ml TGF-β during various time windows (Fig. 4d). Within cells with the highest initial mCitrine-PPARG expression, we observed mCitrine-PPARG downregulation under TGF-β, but only if it was applied at the time of passaging. In contrast, TGF-β application 24h after replating did not lead to a noticeable mCitrine-PPARG downregulation in adipocytes compared to control (Fig. 4e). We concluded that long-term downregulation of adipocyte markers occurs only when TGF-β and mechanical stimuli co-occur.

### The effect of mechanical stress on PPARγ expression is partly mediated by Rho kinase

In order to better characterize the molecular basis of the mechanical stimulus introduced by cell passaging, we took advantage of an apparent transient drop in mCitrine-PPARG expression in adipocytes immediately after cells were passaged (Fig. 4f, Supplementary Fig. S4). We reasoned that if this transient disruption of PPARγ expression was caused by a mechanical stimulus introduced by passaging, then inhibition of the causative mechanotransduction pathway would prevent the mCitrine-PPARG drop. Based on the previously observed dependence of TGF-β-induced PPARγ downregulation on the presence of fibronectin (Fig. 3f), we focused on the mechanotransduction pathways associated with integrin signaling. To this end, we applied several small molecule inhibitors and quantified mCitrine-PPARG levels at 0, 12 and 24 hours after passaging in the subset of cells with the highest initial mCitrine-PPARG levels (Fig. 4f). Rho inhibitor I C3 rescued high mCitrine-PPARG levels while focal adhesion kinase (FAK) and Rho-associated protein kinase (ROCK) inhibitors (PND-1186 and Y27632, respectively) showed no effect (Fig. 4g). In summary, we identified Rho kinase as a possible mediator of mechanical stimuli-driven PPARγ downregulation in adipocytes.

### TGF-β pathway activation is inhibited in adipocytes when a mechanical stimulus is absent

The mechanotransduction-dependent downregulation of PPARγ by TGF-β prompted us to assess the interaction between canonical SMAD-dependent TGF-β signaling and PPARγ. In the TGF-β signaling cascade activation of TGF-β receptor leads to phosphorylation of transcriptional effector proteins SMAD2 and SMAD3 (SMAD2/3) which, together with SMAD4, stimulate TGF-β-dependent transcription. Inhibition of SMAD3 by PPARγ^20,21^ and inhibition of the PPARγ positive feedback partner C\EBPα^22^ by activated SMAD3^23^ were previously described. In support of such a double-negative feedback system between TGF-β signaling and the adipocyte transcriptional network, we observed that co-treatment with the small molecule PPARγ agonist rosiglitazone rescued TGF-β-induced loss of PPARγ expression in primary replated adipocytes (Fig. 5a). To test the efficiency of TGF-β pathway activation as a function of PPARγ expression level, we introduced a live fluorescent reporter of SMAD2/3 transcriptional response (SBE4:mScarlet-I-NLS, Fig. 5b) into the mCitrine-PPARG OP9 cell line. This reporter enables detection of rapid changes in TGF-β signaling-dependent gene expression (Fig. 5c). When a population of differentiated mCitrine-PPARG OP9 cells was treated with TGF-β, there were no observable changes in mCitrine-PPARG expression within 12 hours of live imaging (Fig. 5d-e). During that time upregulation of the TGF-β reporter was restricted to cells with the lowest initial mCitrine-PPARG expression, and co-treatment with rosiglitazone did not lower TGF-β signaling activity in this group (Fig. 5f). The lack of activation of the reporter in mCitrine-PPARG-high cells was not due to insufficient TGF-β concentration as increasing the TGF-β dose 10-fold did not cause TGF-β reporter upregulation in cells with high mCitrine-PPARG expression (Fig. 5g). We concluded that at steady state canonical TGF-β signaling is inhibited in PPARγ-expressing cells. Inhibition of the canonical TGF-β signaling in PPARγ-expressing cells could not be explained by inhibition of SMAD2/3 translocation to the nucleus (Fig. 5h-i), suggesting that it occurred at a later step in the signaling cascade, perhaps through affecting transcriptional activity of SMAD2/3.

**Figure 5.**
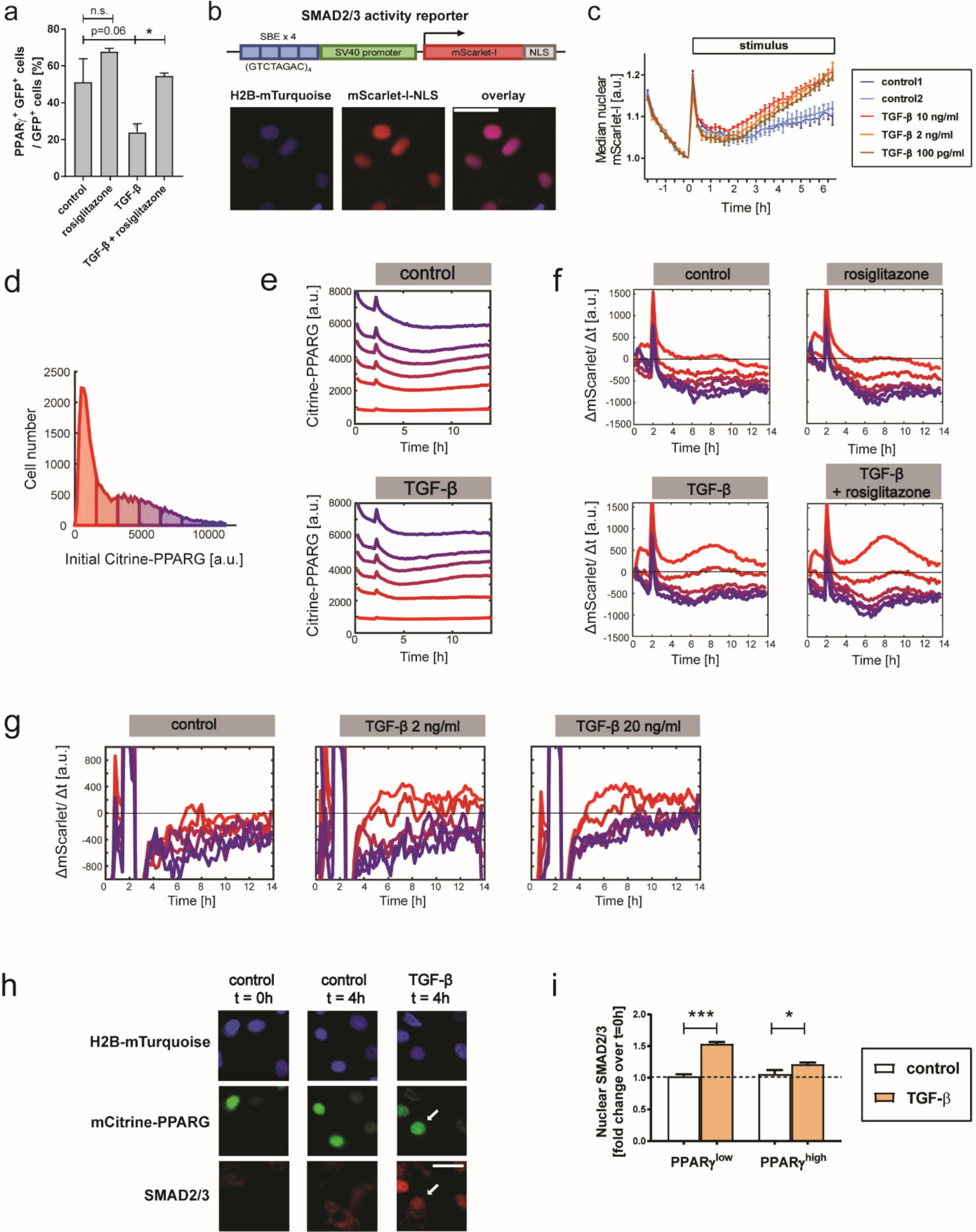
In a population of differentiated OP9 cells TGF-β signaling activation is restricted to PPARγ-low cells. **a** Co-treatment with PPARγ agonist rosiglitazone partially rescues molecular changes associated with TGF-β treatment of SVF-derived adipocytes. Analysis of cells at day six after replating. Stimuli applied continuously beginning at replating. n=3 technical replicates, One-way ANOVA with Sidak correction; *, p<0.05. **b** Schematic of the live fluorescent reporter of SMAD2/3 transcriptional response, SBE4:mScarlet-I-NLS. Representative fluorescent images of undifferentiated OP9 cells are shown. Scale bar: 100 µm. **c** Live fluorescent reporter SBE4:mScarlet-I-NLS allows detection of TGF-β signaling pathway activity. Undifferentiated SBE4:mScarlet-I-NLS OP9 cells were treated with various concentrations of TGF-β after initial 2h of pre-incubation with basal media. For each single cell trace, nuclear signal was normalized by t=0 h. Mean from n=3 technical replicates and S.E.M. are shown. **d** The distribution of single-cell mCitrine-PPARG expression in the last frame before stimulus addition (2h), used to assign cells to bins in panels e-f, shown for the control group. **e** Time course analysis of median PPARγ expression in six bins depending on initial mCitrine-PPARG expression in differentiated non-replated OP9 cells. Data for untreated control and cells treated with 2 ng/ml TGF-β added at 2h are shown. **f** Time course analysis of TGF-β-dependent transcriptional response depending on the initial PPARγ expression (six bins) shows strong reporter upregulation, indicated by positive values of the change in integrated nuclear mScarlet-I signal over time (ΔmScarlet/Δt), in the cell bin with the lowest initial mCitrine-PPARG expression. Median trace for each bin is shown. 12h of treatment with TGF-β (2 ng/ml), rosiglitazone (1 µM), TGF-β and rosiglitazone, or with basal media in control, started after two hours of pre-incubation with basal media. Median mCitrine-PPARG expression traces for each bin are shown. **d-f** Results of one experiment representative for three independent experiments. **g** Increasing TGF-β dose to 20 ng/ml leads to upregulation of SBE4:mScarlet-I-NLS promoter in the same mCitrine-PPARG-low populations as when using 2 ng/ml TGF-β. Consistent changes in fluorescence during the first 2h of experiment attributable to illumination settings. **h** TGF-β treatment leads to efficient SMAD2/3 translocation into the nucleus within 4h both in PPARγ-low and PPARγ-high cells. Differentiated non-replated OP9 cells were treated with TGF-β and subjected to IF staining of SMAD2/3 and PPARγ. Representative fluorescent images prior to TGF-β stimulation and at 4h are shown. Arrows denote a PPARγ-high cell with nuclear SMAD2/3 localization after 4h of TGF-β treatment. Scale bar: 100 µm. **i** Quantification of nuclear SMAD2/3 intensity in PPARγ-high and PPARγ-low cells treated for 4h with TGF-β or control media, normalized to values at t=0h. Two-tailed Student *t* test; *, p<0.05; ***, p<0.001. **h-i** Results of one experiment representative for two independent experiments.

## Discussion

The role of sufficiently high mechanical resistance for myofibroblast maturation from tissue-resident fibroblasts has been previously described^24^. Here, we uncover a novel mechanism by which mechanical inputs such as those introduced by plating adipocytes on a sufficiently stiff fibronectin-containing ECM facilitate early steps of the adipocyte identity loss at the onset of adipocyte-myofibroblast conversion. Through interference with the adipocyte-specific molecular circuitry, activation of mechanotransduction pathways such as Rho kinase pathway allows for full TGF-β signaling activation in adipocytes, which in turn inhibits the adipocyte state and causes a switch to a myofibroblast or other cell states.

We observed a critical role of the transcription factor PPARγ in preventing the loss of adipocyte state. PPARγ expression was lost early during the adipocyte-myofibroblast transition and the PPARγ agonist rosiglitazone prevented this loss. This is in line with known anti-fibrotic effects of thiazolidinediones (TZDs)^20,25^. We observed that high PPARγ expression in a heterogeneous population of differentiated OP9 cells was correlated with the lack of transcriptional response to the profibrotic cytokine TGF-β. The abundance of TGF-β receptors in the cell membrane strongly decreases during fat differentiation^26^ which could potentially explain the decreased TGF-β sensitivity in adipocytes. However, we observed robust translocation of SMAD2/3 into the nucleus irrespective of PPARγ expression level, indicating a block later in the TGF-β signaling cascade, similar to previous observations in lung cancer cells where PPARγ activity inhibits transcriptional activity of SMADs^27^. As TGF-β is a pleiotropic and widespread signaling molecule, direct targeting of the TGF-β pathway in fibrosis treatment is likely to lead to side-effects. Our results suggest that the biological effects of TGF-β in fat fibrosis may be partly counteracted by the reinforcement of the adipocyte molecular circuitry, rather than or in combination with targeting TGF-β signaling directly. Building on these results to elucidate the molecular mechanism by which SMAD-dependent transcription is prevented in cells with high PPARγ expression is an exciting future area of study.

In addition to TGF-β present in culture media, a second stimulus, which was introduced by replating cells at subconfluence, allowed for TGF-β-dependent PPARγ downregulation in adipocytes. Importantly, both stimuli needed to co-occur to obtain long-term PPARγ downregulation. By modulating ECM composition and stiffness, we concluded that interaction with a stiff fibronectin-containing ECM can serve as the second stimulus which, in addition to TGF-β presence, leads to PPARγ downregulation in adipocytes. Importantly, apart from the experiments in which adipocytes were plated on polyacrylamide gels coated with either fibronectin or laminin (Fig. 3d-f), in all other experiments cells were plated on uncoated tissue culture plastic or glass. In these cases, ECM proteins such as fibronectin would likely be present in the serum or produced by the cells themselves since adipogenic cells, especially progenitor cells, are known to produce ECM which contains fibronectin as one of the major components^28^. Also, although we cannot rule out that other ECM proteins may play a role in the observed PPARγ downregulation in adipocytes, we did observe that inhibition of Rho kinase signaling, which is activated downstream of the integrin binding to fibronectin^29^, partially rescues the PPARγ downregulation observed in adipocytes plated on tissue culture glass. A requirement for integration of TGF-β presence and high ECM stiffness has also been reported for chondrocyte differentiation^30^.

High substrate stiffness can increase the amount of biologically active TGF-β through the release of latent TGF-β from ECM under stretch^31,32^ in a fibronectin-dependent mechanism^33^. However, we did not observe an increase in TGF-β-dependent transcription nor PPARγ downregulation in cells with high PPARγ expression when we significantly increased the concentration of TGF-β in the media. This suggests that a simple increase of biologically active TGF-β is not the only mechanism by which high stiffness and fibronectin presence in the ECM contribute to the TGF-β-induced downregulation of PPARγ in adipocytes. ECM stiffness also affected the percentage of adipocytes which showed nuclear translocation of YAP. The threshold of ECM stiffness which was required to allow for TGF-β-dependent PPARγ downregulation and YAP translocation appeared to fall between Young’s modulus of 26 kPa and 90 kPa for primary adipocytes. However, examination of a large number of OP9 adipocytes across a wide range of gel stiffnesses suggested a more linear relationship between ECM stiffness and mechanotransduction in the presence of TGF-β (Supplementary Fig. S5). Fibrotic tissue is much stiffer than most soft tissues (20-100 kPa vs. 0.1 to 1 kPa)^34,35^ and typically shows upregulation of fibronectin and TGF-β1^7^. Taken together, it seems plausible that an existing area of fibrotic tissue adjacent to adipocytes may provide the increase in local ECM stiffness, change in ECM composition and high local TGF-β concentrations that are required to induce adipocyte-myofibroblast transdifferentiation (Fig. 6). In fact, the interface between breast tumors, characterized by high stiffness^36^, and mammary fat contains adipocyte-derived fibroblast-like cells termed adipocyte-derived fibroblasts (ADFs) which exhibit some hallmarks of myofibroblasts^37^. ADFs are a type of cancer-supporting stromal cells, and finding ways to prevent loss of adipocyte identity in this context could be a novel anticancer therapeutic approach.

**Figure 6.**
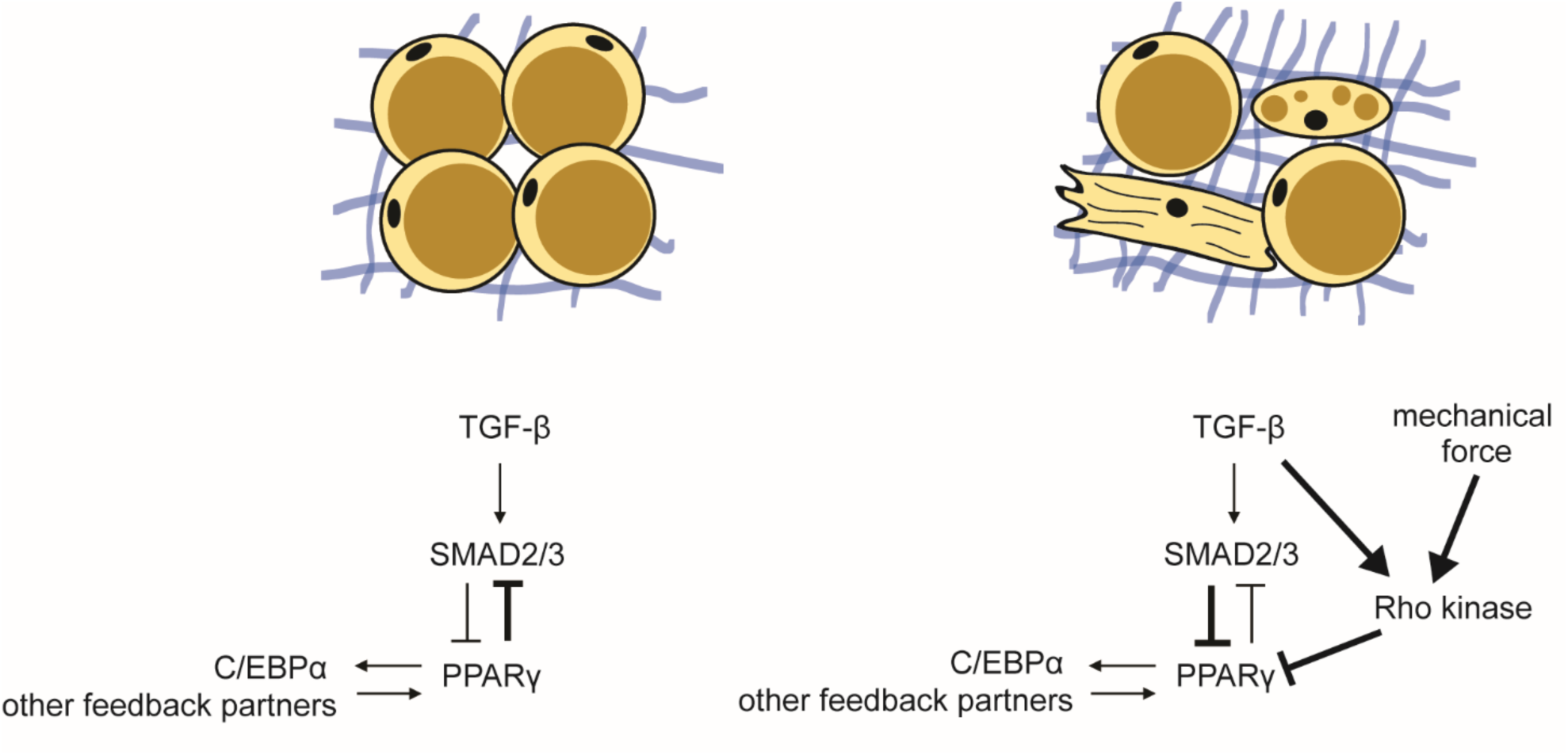
Model of the onset of adipocyte state loss. Activation of mechanotransduction pathways, for example through activation of Rho or YAP, perturbs the adipocyte-specific positive feedback system which includes PPARγ. When a mechanical force is not sensed by an adipocyte this PPARγ feedback system prevents activation of canonical TGF-β signaling.

Our findings on the inhibitory effect of adipocyte state on TGF-β signaling activation may be applicable to more general mechanisms of fibrosis. Certain tissue-specific cell types which constitute the myofibroblast source in fibrosis, such as hepatic stellate cells in liver^38^ and lipogenic fibroblasts in lung^39^, share certain molecular characteristics with adipocytes including PPARγ expression^39,40^. Understanding the precise mechanisms by which mechanical inputs and TGF-β counteract PPARγ activity at the onset of fibrosis could aid in finding therapeutic antifibrotic targets, for example through manipulating mechanotransduction signaling pathways. In fact, a focal adhesion kinase inhibitor has been successfully used in preventing fibrosis development in a mouse model^41^. However, it is not clear at which point in the multi-step process of fibrosis the FAK inhibition acts. Here we identified Rho kinase as a possible target whose inhibition can counteract PPARγ downregulation in adipocytes after mechanical stimulation associated with plating on a stiff substrate, but it needs to be determined whether these pathways are directly responsible for the mechanical response in adipocytes. It also remains to be tested whether Rho kinase enables TGF-β signaling transduction after interaction with a stiff fibronectin-containing ECM.

In this study we identify a molecular mechanism which prevents loss of adipocyte identity. Our data support the role of mechanical stimuli as a possible second stimulus which allows TGF-β to downregulate adipocyte-specific transcriptional program at the onset of adipocyte-myofibroblast transition. Our findings underscore the need for further investigation into the role of ECM properties in regulating adipocyte behavior in the context of fibrosis.

## Methods

### Animals

All animal studies were conducted according to Stanford University guidelines. Mice were purchased from Jackson Laboratory. mT/mG B6.129(Cg)-Gt(ROSA)26Sortm4(ACTB-tdTomato,-EGFP)Luo/J (cat. 007676) and nT/nG B6;129S6-Gt(ROSA)26Sortm1(CAG-tdTomato*,-EGFP*)Ees/J (cat. 023035) mice were bred to B6;FVB-Tg(Adipoq-cre)1Evdr/J mice (cat. 010803).

### Cell culture and differentiation

Stromal vascular fraction (SVF) was isolated from inguinal subcutaneous fat pads of 4-8-week old female and male mice using a previously published approach^42^. Fat pads were minced and digested in a solution of collagenase type D (Roche, 11088866001, 1 mg/ml) and Dispase II (Sigma-Aldrich, D4693, 1 mg/ml) in PBS with 1mM CaCl_2_ for 40 minutes at 37°C with shaking. The digest was passed through sterile nylon mesh and centrifuged at 300 RCF for 5 minutes. The SVF-containing pellet was resuspended in culture medium (DMEM with 10% FBS + 100U/mL pen/strep) with 2.5 mg/ml amphotericin B for 2-4 hours, which was then replaced. Cells were grown in the presence of 2.5 mg/ml amphotericin B for up to seven days before the start of differentiation protocol. To differentiate the SVF, cells were plated in 12-well cell culture plates at 120,000 cells per well (day -1). At day 0, cells were treated with 250mM IBMX (Sigma-Aldrich), 1mM dexamethasone (Sigma-Aldrich), 1.75nM insulin (Sigma-Aldrich) and 500 nM rosiglitazone (Cayman Chemical) in culture medium. At day 2, cells were treated with 1.75nM insulin and 500nM rosiglitazone in culture medium, and at day 4 with 1.75nM insulin in culture medium for two more days. Differentiated SVF cells were maintained in culture medium with 1.75nM insulin afterwards.

OP9 cells were cultured in MEM-α media (Invitrogen) containing 100 units/mL Penicillin, 100mg/mL Streptomycin, and 292 mg/mL L-glutamate. The base media also contained either 20% Fetal Bovine Serum (FBS) for cell expansion or 10% FBS for cell differentiation. To induce differentiation of OP9 cells, a standard DMI protocol was used: confluent cells were treated with 250 mM IBMX (Sigma-Aldrich), 1 mM dexamethasone (Sigma-Aldrich), and 1.75nM insulin (Sigma-Aldrich) for 48h, followed by 1.75nM insulin for 48h. Afterwards differentiated OP9 cells were maintained in differentiation medium with 1.75nM insulin.

Mouse TGF-β 1 recombinant protein was obtained from Affymetrix (#14-8342-62) and used at the concentration of 2 ng/ml unless stated otherwise. The following chemical inhibitors were used: Y27632 (Fisher Scientific, 10µM), PND-1186 (VS-4718, Fisher Scientific, 1µM), and Rho Inhibitor I C3 (Cytoskeleton, 0.5 µg/ml).

### Generation of SBE4:mScarlet-I-NLS reporter OP9 line

For the cloning of fluorescent reporter of TGF-β transcriptional response (SBE4:mScarlet-I-NLS), Gibson Assembly Master Mix (New England Biolabs) was used according to manufacturer’s protocol. PiggyBac vector PB-CMV-MCS-EF1a-Puro (System Biosciences) was first modified to include blasticidin resistance gene instead of the puromycin one and linearized using SfiI and XbaI. Smad2/3 response element was amplified from SBE4-Luc construct, which was a gift from Bert Vogelstein (Addgene plasmid # 16495)^43^, and cloned upstream of mScarlet-I sequence^44^ with inframe nuclear localization sequence (NLS). The construct was introduced into mCitrine-PPARG H2B-mTurquoise OP9 cells by co-transfection with PiggyBac transposase vector, followed by selection with blasticidin (Thermo Fisher Scientific).

### RT-qPCR

Gene expression was quantified using Go Taq Green Master Mix (Promega) and the LightCycler 480 Instrument II (Roche). Primers are listed in Supplementary Table 1. Expression of the cyclophilin gene was used for normalization using the ΔΔCt method.

### Polyacrylamide gel preparation

Polyacrylamide gels were prepared in 12-well glass-bottom plates (Cellvis, P12-1.5H-N), which were activated through consecutive incubation with 2% 3-aminopropyltrimethoxysilane (Acros Organics) in isopropanol for 15 min at room temperature, three washes with water and incubation with 2.5% glutaraldehyde in water for 30 min at room temperature. Finally, the plates were washed three times with water and dried. To prepare the gels, round glass coverslips were plasma cleaned and incubated with 50 ug/ml solution of human fibronectin (Corning) or mouse laminin (Sigma-Aldrich) in PBS. Polyacrylamide gels of ∼200 µm thickness were prepared by mixing 2× acrylamide / bisacrylamide stock with PBS, de-gassing the mixture, addition of 0.5% volume of 10% ammonium persulfate (VWR) in water and 0.2% tetramethylethylenediamine (TEMED, Fisher Scientific). The solution was then pipetted onto the 12-well plates and covered with protein-coated coverslips. The gels were allowed to polymerize overnight followed by three washes with PBS prior to use. The 2X acrylamide / bisacrylamide stocks were prepared based on published recipes for the required stiffnesses^45^ and used throughout the study.

### Immunofluorescence (IF) staining

To minimize cell loss due to detachment, cells grown on polyacrylamide gels were pre-fixed by the addition of paraformaldehyde (PFA) to the final concentration of 4% directly into the growth media and incubation for 10 min. All cultured cells were fixed with 4% PFA in PBS for 30 min at room temperature, followed by three washes with PBS. Cells were then permeabilized with 0.1% Triton X-100 in PBS for 15 minutes on ice, followed by blocking with 5% bovine serum albumin (BSA, Sigma Aldrich) in PBS. The cells were incubated with primary antibodies in 2% BSA in PBS overnight at 4°C: mouse anti-PPARγ (Santa Cruz Biotech, sc-7273, 1:1,000), rabbit anti-CEBPα (Santa Cruz Biotech, sc-61, 1:1,000), mouse anti-YAP (Santa Cruz Biotech, sc-101199, 1:500), chicken anti-GFP (Fisher Scientific, NB1001614, 1:1,000), rabbit anti-α-SMA (Abcam, ab5694, 1:500). After washing, cells were incubated with Hoechst (1:10,000) and secondary antibodies in 2% BSA / PBS overnight at 4°C. Secondary antibodies included AlexaFluor-conjugated anti-rabbit and anti-mouse antibodies (1:1000, Invitrogen) and anti-chicken AlexaFluor488 antibody (Thermo Fisher Scientific, A11039, 1:1,000). Where indicated, lipids were co-stained by adding BODIPY 493/503 (1mg/ml, Molecular Probes #D-3922) to secondary antibody solution. Cells were washed three times with PBS prior to imaging.

### Fluorescent imaging

Imaging was conducted using either an ImageXpress MicroXL (Molecular Devices, USA) or a 3i (Nikon) epifluorescent microscope with a 10X objective, with the exception of fixed cells seeded on polyacrylamide gels, which were imaged using 20X objective. Live fluorescent imaging was conducted at 37°C with 5% CO_2_. A camera bin of 2×2 was used for live imaging and 1×1 was used for fixed imaging. Cells were plated in optically clear 96-well plates: plastic-bottom plates (Costar, #3904) for fixed imaging or glass-bottom µ-Plate (Ibidi, #89626) for live imaging. Living cells were imaged in FluoroBrite DMEM media (Invitrogen) with 10% FBS, 1% Penicillin/Streptomycin and insulin to reduce background fluorescence. Depending on the experiment, images were taken every 12-15 min in different fluorescent channels: CFP, YFP and/or RFP. Total light exposure time was kept less than 500 ms for each time point. Several non-overlapping sites in each well were imaged. Cell culture media were changed at least every 48h.

To detect myofibroblasts, cells were fixed and subjected to IF staining against α-SMA as described. Following the staining, the same sites were re-imaged, and cells were incubated in solution of Alexa Fluor 647 phalloidin (Cell Signaling, 1:1,000 in PBS) for 30 min at room temperature to stain total actin. After 3 × PBS washes, the same sites were re-imaged. Total actin staining was used for cell segmentation and cell morphology measurement, α-SMA staining was used to quantify α-SMA and stress fibers.

### Imaging data processing

With the exception of quantification of YAP translocation, data processing of fluorescent images was conducted in MATLAB R2016a (MathWorks). YAP translocation was quantified using manual categorization of cell images using ImageJ in blinded experiments. Unless stated otherwise, fluorescent imaging data were obtained by automated image segmentation, tracking and measurement using the MACKtrack package for MATLAB^46^. Quantification of PPARγ- and C\EBPα-positive cells was based on quantification of mean fluorescence signal over nuclei. Cells were scored as PPARγ-and C/EBPα-positive if the marker expression level was above a preset cut-off determined by the bimodal expression at the earliest analyzed time point. GFP-positive cells were scored based on the mean value of GFP fluorescence signal measured over cell nucleus being above a preset cutoff determined by analysis of the distribution in the population.

In myofibroblast phenotype detection experiments, imaging sites which were out of focus were removed from analysis, and cells were initially filtered based on integrated Hoechst signal (nucleus size), cell area size (to remove incorrectly segmented cells) and mean nuclear GFP fluorescence (to filter for GFP-positive cells). The methodology for actin fiber measurement was based on published approaches^47,48^. The algorithm to quantify actin score was added to the MACKtrack package (actin module). Automated myofibroblast detection was optimized using Classification Learner app and a linear support vector machine (SVM) classifier. Out of six initial variables (actin score, integrated actin score, cell area, mean cellular α-SMA expression, integrated cellular α-SMA expression, axis ratio) used to train the model on a training dataset, actin score and mean cellular α-SMA expression were chosen as producing maximal accuracy of myofibroblast phenotype prediction compared to manual scoring. In subsequent experiments, cut-off values for actin score and mean cellular α-SMA expression were chosen arbitrarily based on value distribution in control group and were used consistently for all time points and conditions tested.

For the quantification of cell area of SVF-derived cells, images in the GFP channel were processed using ImageJ 1.52h by auto thresholding using the IsoData method and measurement of cell area. Mean nuclear PPARγ expression in the same GFP-positive cells was quantified automatically using the MACKtrack package.

For live imaging data of OP9 cells, CFP channel was used for nuclear segmentation and cell tracking. Obtained single-cell traces were filtered to remove cells absent at endpoint, traces with more than 10 empty frames and a fraction of traces with maximal changes of PPARγ intensity, quantified as the maximum of a moving integral of the squared difference between PPARγ intensity and local average over a window of double the length of the window used for cell tracking. The filtering for the changes of PPARγ intensity was according to a set cut-off for all conditions, and the cut-off was chosen so that only up to 2% of traces were removed in control.

If cells were binned according to their PPARγ expression, cells were binned based on their mean nuclear PPARγ expression in the first frame of the experiment, with the exception of SBE4:mScarlet-I-NLS reporter cells, which were binned based on the mCitrine-PPARG expression in the last frame prior to addition of stimulus.

To quantify activity of the SBE4:mScarlet-I-NLS reporter, mScarlet-I signal was measured integrated over the whole nuclear area and recalculated as the change over the preceding frame.

Median of single-cell Δ[Integrated mScarlet-I-NLS]/Δt traces was then smoothened using a moving average over time window equal to double the number of frames used for accurate single-cell tracking in MACKtrack. If a cell trajectory present at the beginning of the experiment (parent cell) split into more trajectories (daughter cells), the mScarlet-I signal values for the parent were calculated as the mean of daughter cell trajectories.

### Statistics

Unless specified otherwise, data are expressed as mean +/-standard error of the mean (S.E.M). p-values < 0.05 were considered statistically significant. Analyses were performed using PRISM software v. 7.04.

## Data availability

All relevant data from this manuscript are available upon request.

## Acknowledgements

This work was supported by National Institutes of Health RO1-DK101743, RO1-DK106241, P50-GM107615, a Stanford BioX Seed Grant, and a Stanford Diabetes Research Center Seed Grant (to M.N.T.), American Heart Association Postdoctoral Fellowship 18POST34030448 and Stanford Center for Systems Biology Seed Grant (to E.B.M.), NIH F32 Postdoctoral Fellowship 5F32DK114981-02 (to B.T.), and NIH F31 Predoctoral Fellowship 1F31DK112570-01A1 (to M.L.Z.). The authors would like to thank members of Teruel lab for helpful discussions.

## Author contributions

E.B.M. and M.N.T. conceived of the study, E.B.M., C.M., M.L.Z., A.R.D., and M.N.T. designed the experiments. Experiments were conducted by E.B.M., C.M. (polyacrylamide gel preparation), A.S. (RT-qPCR) and M.L.Z. (a subset of live fluorescent imaging experiments). B.T. created scripts for actin fiber measurement and basic MATLAB data filtering. Z.B.N. provided the mCitrine-PPARγ OP9 cell line. E.B.M. analyzed the data. E.B.M. and M.N.T. wrote the manuscript with inputs from all authors.

## Materials & Correspondence

Requests for materials and correspondence should be directed to Ewa Bielczyk-Maczyńska or Mary N. Teruel.

## Supplementary Information

### Supplementary Figures

**Supplementary Figure S1.**
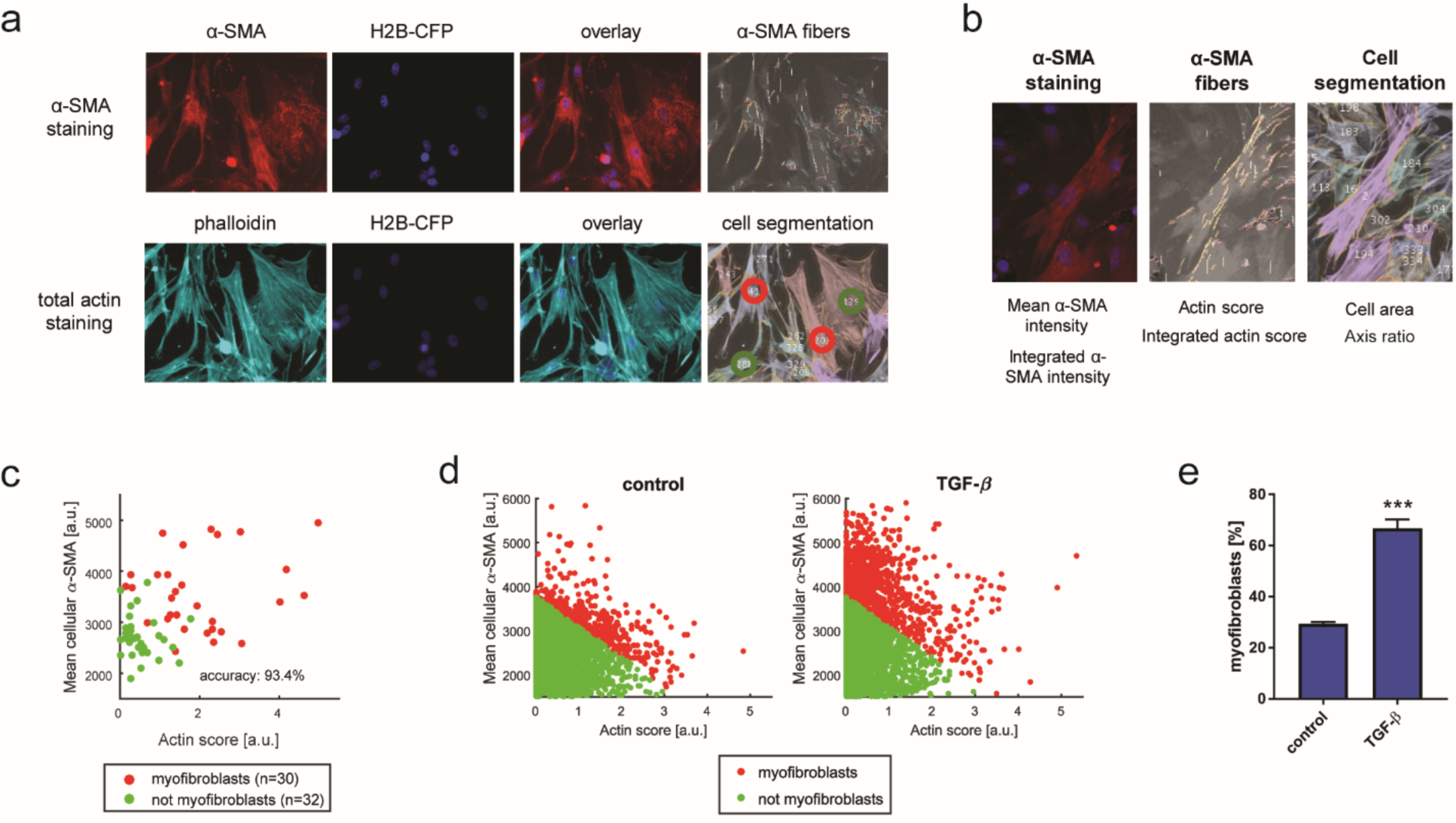
Automated discovery of myofibroblast phenotype using IF staining against α-SMA and total actin. **a** Representative fluorescent images of α-SMA staining and consecutive total actin staining with phalloidin, automated α-SMA fiber detection and cell segmentation of OP9 cells. Cells used for the training dataset were differentiated, replated at subconfluence, and TGF-β was applied 48h later for 48h. A subset of the cells were subjectively categorized as positive (red) or negative (green) for myofibroblast phenotype. **b** Six variables used to optimize automated myofibroblast detection. Actin score represents the maximal fraction of cell area which contains α-SMA fibers oriented in the same direction (directions are color-coded). Integrated actin score represents the sum of actin scores for all directions. Axis ratio represents cell length divided by cell width. **c** Scatter plots of the separation of the myofibroblasts and non-myofibroblasts in the training dataset. Linear support vector machine classifier, based on actin score and mean cellular α-SMA, was chosen based on its maximal accuracy on the training dataset. **d** Scatter plots of the separation of the myofibroblasts and non-myofibroblasts in the validation dataset. Once trained, the classifier was applied to all cells in two other conditions from the same experiment: control and TGF-β-treated from 24h post-replating onwards. **e** Quantification of automatically detected myofibroblasts. n=4 technical replicates. ***, p<0.001.

**Supplementary Figure S2.**
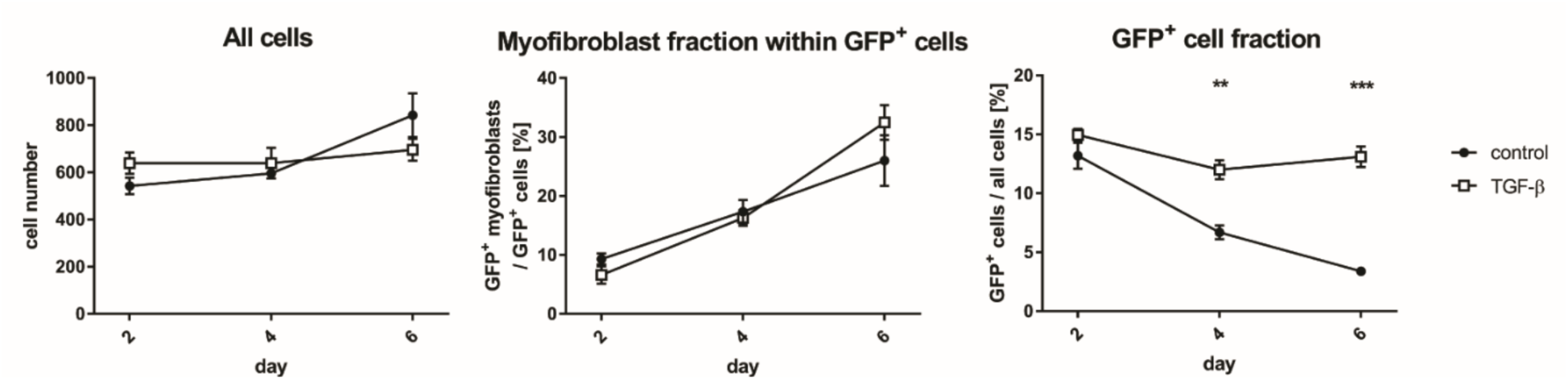
Increase in the number of adipocyte-derived myofibroblasts may be caused by enhanced survival of GFP-positive cells under TGF-β compared to control. Data corresponding to Fig. 1g.

**Supplementary Figure S3.**
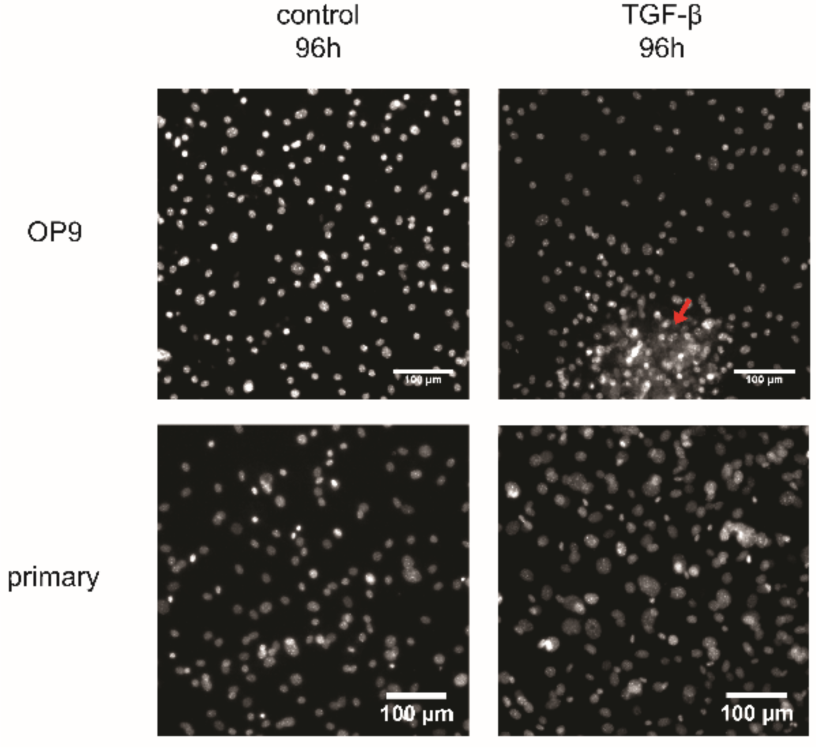
TGF-β treatment leads to clump formation in differentiated OP9 but not SVF-derived primary cells. Differentiated OP9 and SVF were stimulated with control media or TGF-β (2ng/ml) for 96h. Nuclei stained with DAPI are shown. A cell clump present in OP9 cells stimulated with TGF-β is indicated with the red arrow.

**Supplementary Figure S4.**
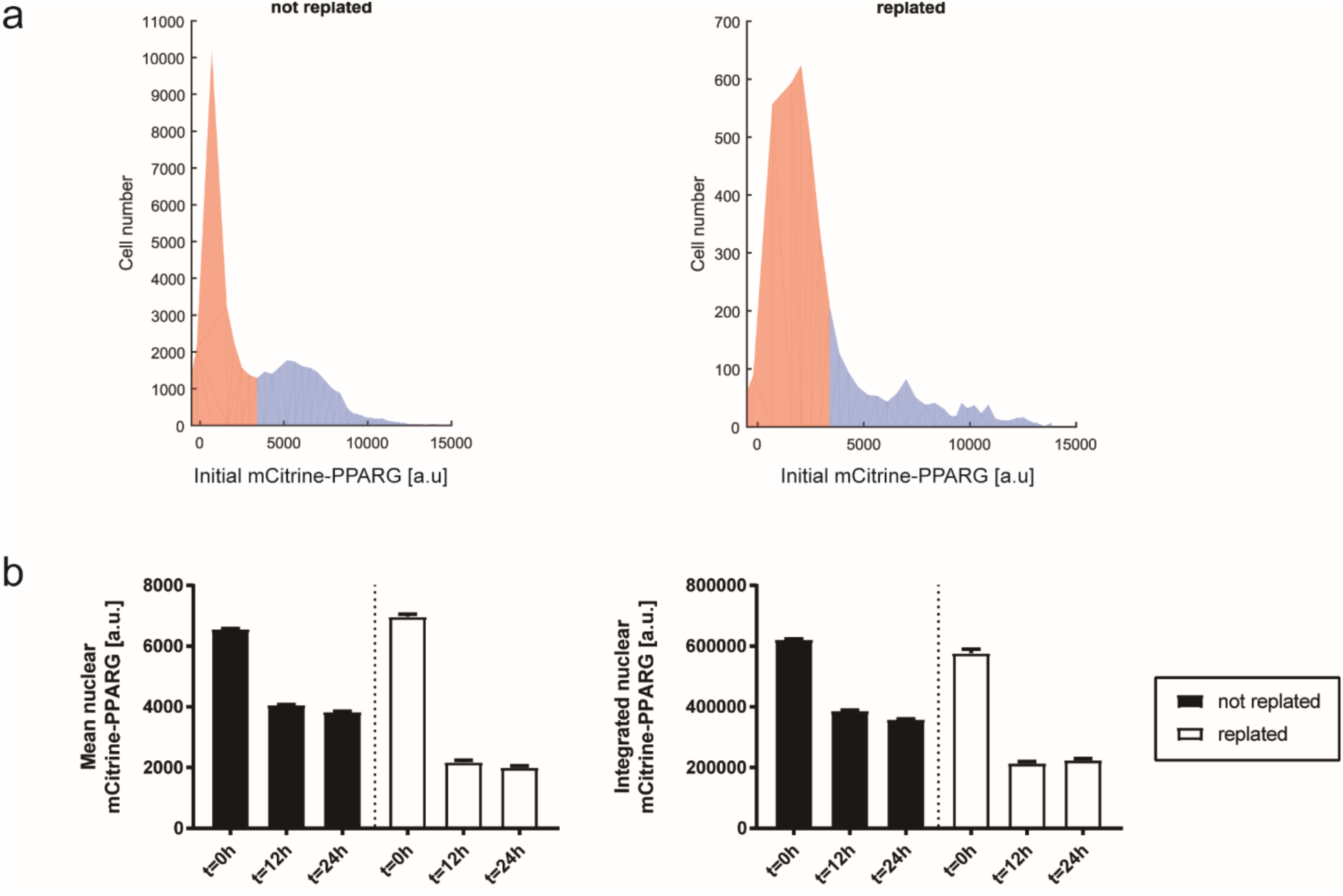
Comparison of mCitrine-PPARG expression in not replated and replated control OP9 cells. Not replated condition used to control for changes in background fluorescence over time. Signal integrated over nucleus used to control for changes in nuclear shape after replating, compared to mean nuclear signal (integrated signal divided by nucleus area). Media changed at t=0h in both conditions. **a** Histograms of mean nuclear mCitrine-PPARG in replated and not replated cells at t=0h. Blue denotes the mCitrine-PPARG-high population analyzed in later steps. **b** Quantification of the mean nuclear and integrated nuclear mCitrine-PPARG signal at 0h, 12h, 24h. Average and S.E.M. for individual traces are shown. Not replated cells: n= 19,723 traces; replated cells: n= 1,259 traces.

**Supplementary Figure S5.**
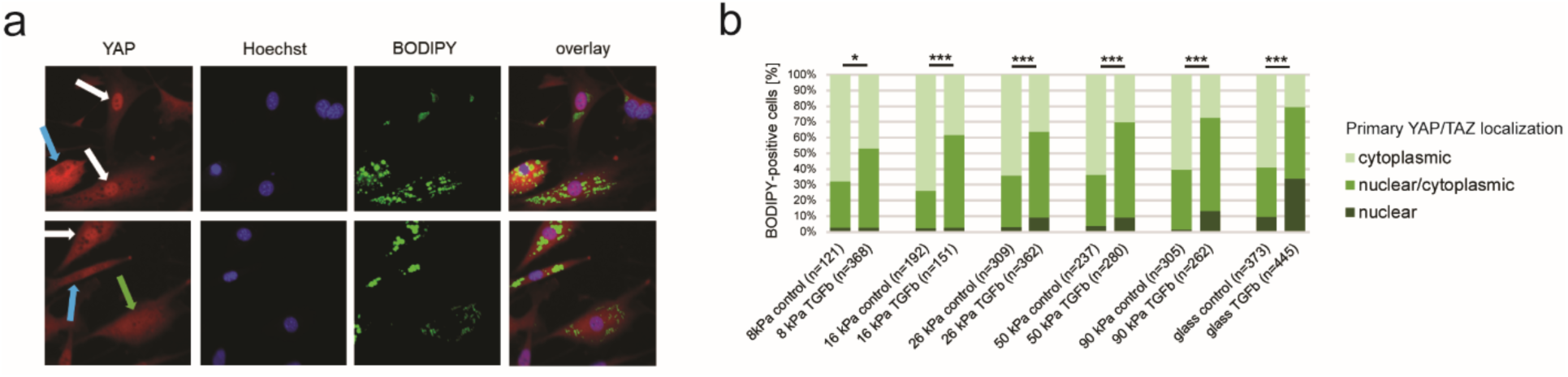
Quantification of YAP localization in a population of differentiated non-transgenic OP9 cells which were fixed 24h after replating on polyacrylamide gels of varied stiffnesses, using the presence of lipid vesicles as a proxy marker for adipocyte state. **a** Representative fluorescent images of two sites, OP9 cells were replated on cell culture plastic. YAP was stained using IF staining and lipids were visualized using BODIPY. Nuclei were counterstained using Hoechst. Cells with predominantly nuclear (white arrows), predominantly cytoplasmic (blue arrows) and no predominant localization (green arrow) of YAP are shown. **b** Quantification of predominant YAP localization in BODIPY-positive OP9 cells depending of the stiffness of fibronectin-covered polyacrylamide gel. Kruskal-Wallis test with Dunn’s multiple comparison correction; *, p<0.05, ***, p<0.001

## Supplementary Tables

**Supplementary Table 1.**
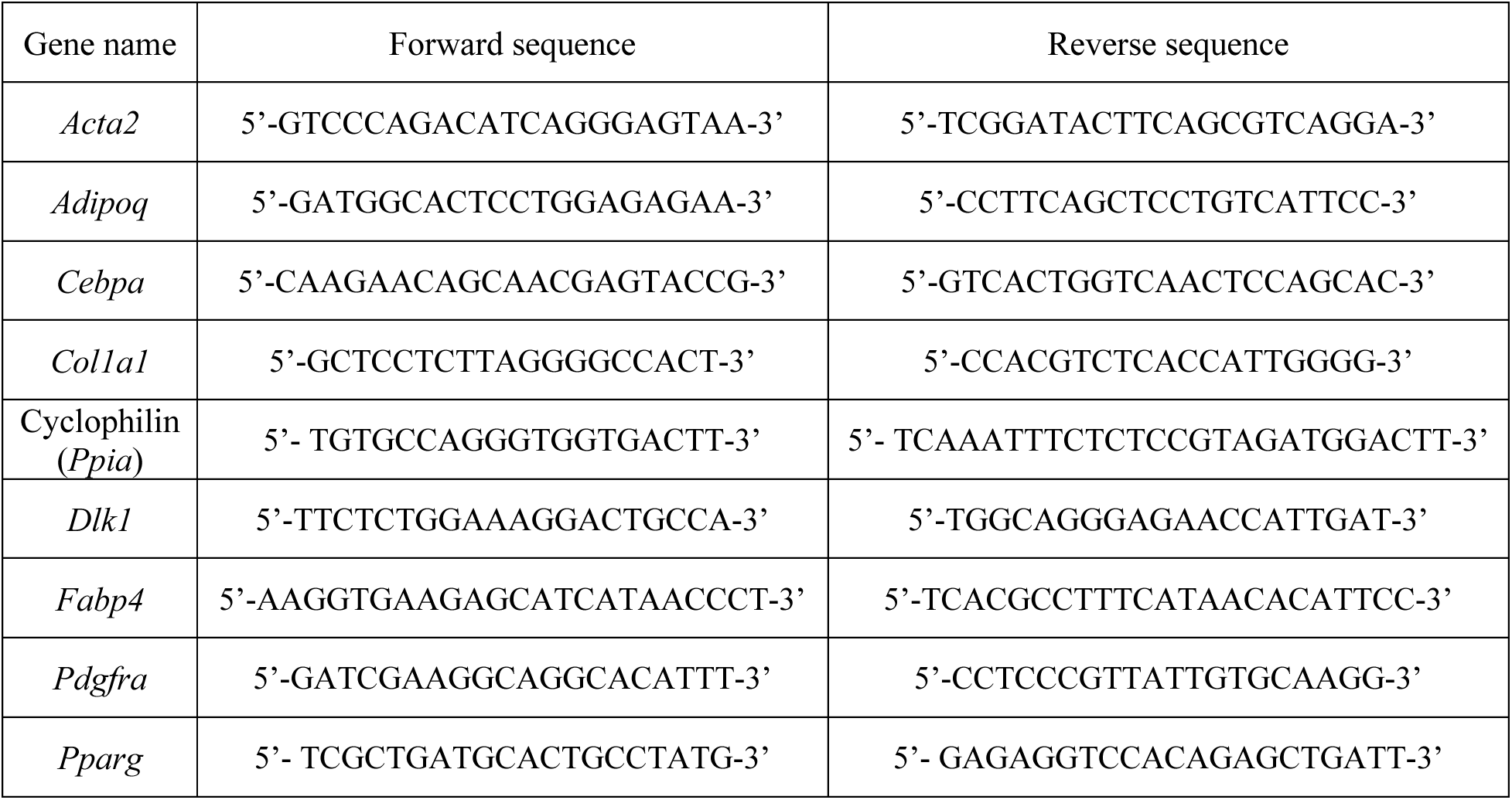
Sequences of RT-qPCR primers used in the study.

